# Forecasts for 80% of wild populations and communities predict no change, and why

**DOI:** 10.1101/2025.06.13.659576

**Authors:** Claudio Bozzuto, Anthony R. Ives

## Abstract

Many retrospective analyses of biodiversity trends reveal a complex picture, including significant declines but also increases. However, prospective analyses (i.e. what near-term trends to expect) remain comparatively rare. Here, we generate and assess forecasts for approximately 43,000 population-level and 10,000 community-level time series from localities across the globe and covering taxa across the Tree of Life. Using unobserved components models and model selection, we found that only 23.5% of wild populations exhibit forecasted time trends (with increasing and decreasing trends roughly equal), while 76.5% show a forecasted constant mean. For communities (temporal *α*- and *β*-diversity), this figure rises to 84.0%. We identified high variation in the rate of change, var(RoC), as the most important predictor of a forecasted constant mean, and it scaled more strongly with intrinsic ecological dynamics than observation noise. Furthermore, biological structure (taxonomy, life-history traits, and locality) contributed to the identification of trends, highlighting that biological factors underlie, in part, predictions of trends. In summary, despite time trends being identifiable in ecological time-series data, parsimonious statistical models will very often forecast no change. Our results suggest that the absence of directional change may be the norm rather than the exception across the Tree of Life.

## Introduction

Since E. O. Wilson’s book “The Diversity of Life” [1] – published the same year as the United Nations “Earth Summit” held in Rio de Janeiro in 1992 – a growing body of research has chronicled an accelerating biodiversity crisis (e.g. [2], and references therein). Numerous studies document alarming shifts in species’ abundance and genetic diversity, in community composition, and in extinction rates. However, whether these trends amount to an impending or unfolding “sixth mass extinction” is contested. For example, Wiens and Saban [3] show that, although contemporary losses are elevated relative to longterm baselines, they lack the catastrophic severity characteristic of past mass-extinction events.

Retrospective temporal analyses of recent biodiversity changes have shown that some taxonomic groups in particular have experienced large declines: examples come from birds (e.g. [4]), fishes (e.g. [5]) and insects (e.g. [6]). Broader studies, on the other hand, have found that the picture of biodiversity trends is more nuanced than a narrative of a pervasive decline implies, with no net change at local scales, and with declining and increasing trends roughly balanced ([7], and references therein). At the same time, dozens of databases have been designed and made publicly available that contain many thousands of time series at the population and community levels (also referred to as assemblages), spanning a wide range of taxa and localities globally (e.g. [8], and references therein). These data collections offer an unprecedented opportunity to investigate biodiversity changes at different temporal and spatial scales. Partly based on these databases, several authors have explored why reported biodiversity trends diverge so widely. One reason involves the diversity of definitions, methods, and approaches adopted [9]. A second reason is methodological, such as the common neglect of correlation structures in the data [8] and the presence of mathematical biases [10] when computing summary trends. A third reason involves heterogeneity among the time-series datasets. Although an impressive volume of records is now available for studies of biodiversity change, these data vary greatly in their temporal extent, reliability, and the types of biodiversity attributes they measure (e.g. [10]).

Prospective studies on expected biodiversity trends primarily focus on capturing the effect of human-caused drivers, including climate change, on future population and community states (e.g. [11,12]). These prospective studies often rely on long-term scenario-based projections of environmental changes, for example projected climate change responses over the next several decades. Studies indicate that if current trends in human-caused stressors continue or worsen in the absence of significant conservation interventions, many systems might experience local biodiversity declines on the order of 10–20% over the next few decades (e.g. [13,14]). In contrast to long-term projections, several authors have recently stressed the importance of near-term forecasts (i.e. a forecast horizon of at most a decade), not least to match the typical timescale of decision-making in biodiversity conservation management [12,15,16]. But despite this change of timescale to better and more actionably understand expected biodiversity trends, developing process-based approaches encompassing mechanistic links between stressors, populations, and ecosystems remains a formidable task. We agree with Lasky et al. [17] who question the usefulness to include processes from a multitude of fields (such as ecophysiology, evolutionary biology, demography): “We argue that answering this critical question is ultimately an empirical exercise requiring a substantial amount of data that have not been integrated for any system to date.”

The number of studies making and reporting near-term statistical forecasts is rapidly growing [18]. Broad assessments across localities, realms and taxonomic groups, on the other hand, continue to be rare. Compared to retrospective biodiversity trend analyses, the currently available data collections have not yet been used extensively to construct a coarse forecasting baseline against which more refined and system-specific approaches can be evaluated. Some of the studies focusing on forecasting have analyzed single taxonomic groups or realms (e.g. [19,20]). Other studies have used several data collections to either assess the performance of forecasting models [8,21] or to investigate the concept of predictability in ecology and conservation [22,23]. Predictability-related methods, however, are mainly insightful when analyzing time series spanning at least 20 years [23]. A considerable fraction of currently available data collections, unfortunately, consists of (very) short time series. For example, in the widely used BioTIME database, only about 50% of the time series span more than seven years, and some only two years [24]. We can expect that adapting approaches from climate change science will significantly improve our ability to predict expected biodiversity changes at different spatial and temporal scales using mechanistic models [12]. But for the time being, we need to answer a simple question: What do parsimonious, robust statistical models show about expected near-term biodiversity trends from typical ecological time series?

For this study we analyzed approximately 43,403 population-level and 10,409 community-level time series from 64 taxonomic classes nested within 6 kingdoms, with time series from throughout the world. We found that for the majority of populations and communities, parsimonious statistical models show a forecasted constant mean rather than a directional time trend. We then investigate important drivers of this result to determine what are the characteristics of typical ecological time-series data that are associated with a forecast of no change. Our findings do not argue against the possibility of future biodiversity loss. They do, however, challenge the prevailing narrative that past trends are expected to “automatically” continue. We show that, based on data from high-profile databases, the most frequent parsimonious forecast is for no expected change, even though trends may be visible in the data.

## Materials and methods

### Time-series data

We used time-series data from ten different sources and databases (see ESM: Table S1 for additional details, including access information): BioTIME [24], GBIF [25], GPDD [26], LPI [27], LTER [28], MASTREE+ [29], RivFishTIME [30], TREAM [31], Swiss long-term phytoplankton data [32], and Swiss long-term macrofungal data [33]. To avoid the potential repeated inclusion of the same time series, we used each data collection for a specific taxonomic group and/or realm. Moreover, where possible we used a data collection for either population-level or community-level analyses, not both. We achieved an almost global coverage for population-level data (ESM: Figure S1; community-level: ESM: Figure S2), and a broad population-level taxonomic coverage with 64 classes nested within six kingdoms (ESM: Figure S3; community-level: ESM: Figure S4).

We analyzed time series spanning at least 5 years. We allowed a proportion of missing values of at most 25% (of which at most 50% occurred consecutively), and we excluded binary data (e.g. presence-absence time series). We required the first and last time series value to be present (not missing) and non-zero [10]. Unless time series had already been transformed and contained negative values, we transformed the data using the inverse hyperbolic sine function. This transformation resembles the log-transformation for positive values, but can handle zeros. Finally, we z-transformed all resulting time series. In ESM: Section S1 we provide additional details on data preprocessing.

At the community-level we considered two biodiversity metrics covering temporal *α*-diversity and temporal *β*-diversity. To construct *α*-diversity time series, we computed year-specific Hill numbers [34]. In particular, we used the _1_*D* metric which is the exponential of the Shannon entropy. To construct *β*-diversity time series, we compared each pair of temporally consecutive community compositions and quantified their similarity using Ruzicka’s metric [35]. This metric is related to Jaccard’s similarity metric but additionally accounts for population abundances instead of identities alone. We then transformed these time series for further analysis as described above for populations. To address the potential confounding effect of changes through time in sampling effort, which can affect measures of diversity, we conducted a proof-of-concept sensitivity analysis using the BioTIME database (ESM: Section S1). For this analysis, we used total community abundance as a proxy for sampling effort. Community time series whose reported abundance showed a trend (i.e. were best described by a model with a forecasted time trend) were temporarily excluded, and the resulting proportion of communities with a forecasted constant mean in *α*-diversity was recomputed. This proportion was even higher than in the full dataset, suggesting that our main results are conservative with respect to potential confounding effects of sampling effort.

### Unobserved components models and model selection

Unobserved components models (UCMs) are univariate state-space models which are widely used in many fields of science, including ecology (e.g. [36–38]). They are flexible, explicitly separate measurement error from process error, can handle missing values, and accommodate nonstationary dynamics without the strict assumption of stationarity that is common in other time-series frameworks. As a template for our four UCMs, we used the local linear trend model given by

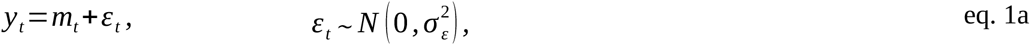

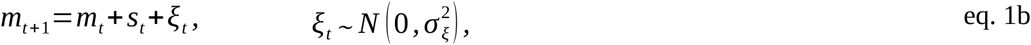

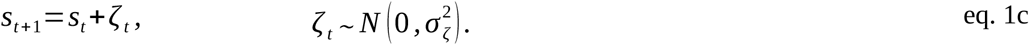

Eq. 1a relates the unobserved true mean *m*_*t*_ to the data, *y*_*t*_, with measurement error. The true mean (eq. 1b) evolves as a random walk and drifts due to a slope which also evolves as a random walk (eq. 1c). Depending on the variances included, many different UCMs can be constructed based on the local linear trend model. Preliminary investigations showed that the full local linear trend model (eq. 1a-c) almost never was selected as the best-fitting model for our collection of time series. Thus, for our analysis we selected the following four models.

- *Deterministic constant*: This is the baseline model, with only eq. 1a included and where *m*_*t*_ *≡m* is a constant. For this model, a single variance has to be estimated.
- *Deterministic trend*: This model includes both the measurement equation (eq. 1a) and the mean equation (1b), which however evolves deterministically. Therefore, *s*_*t*_ ≡ *s* and 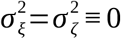, so that only one variance is estimated (eq. 1a).
- *Local level*: This model includes both the measurement equation (eq. 1a) and the mean equation (1b). The mean evolves stochastically as a random walk, but there is no stochastically changing slope. Therefore, 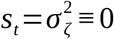, and two variances are estimated (eq. 1a-b).
- *Smooth trend*: This model includes all three equations (eq. 1a-c). However, the mean equation has no stochastic variation of its own. Therefore, 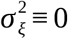, and two variances are estimated (eq. 1a, 1c).

For each population or community time series, we fit all four models and selected the best-fitting one using AIC corrected for small sample size (AICc).

Two of our models produce forecasts with no time trend (the deterministic constant model and the local level model), whereas the other two models allow the forecasted mean to include a linear time trend. Regardless of whether the forecasted mean is constant or changes linearly, fitted UCMs can show a stochastic trend over the timespan of the data. To illustrate this, we analyzed the Eastern monarch butterfly population (Figure 1; ESM: Section S1) (e.g. [39]). Each row shows a selected period of the whole time series (Figure 1a-b), and for each timespan we show all four model fits and include ΔAICc values, where a value of zero designates the best-fitting model for the given time span. The left column in Figure 1 shows the two models with a forecasted constant mean, whereas the right column shows the two models with forecasted linear trends. We highlight three points. First, for the full time series (Figure 1a-b) and the one up to year 2018 (Figure 1c-d), the local level model is the best-fitting one. Here, the mean evolves stochastically and follows the data, including changes in trends (2013, 2018). However, because we can have no knowledge about future stochastic deviations (eq. 1b), the logical and robust consequence is a forecasted constant mean.

**Figure 1.**
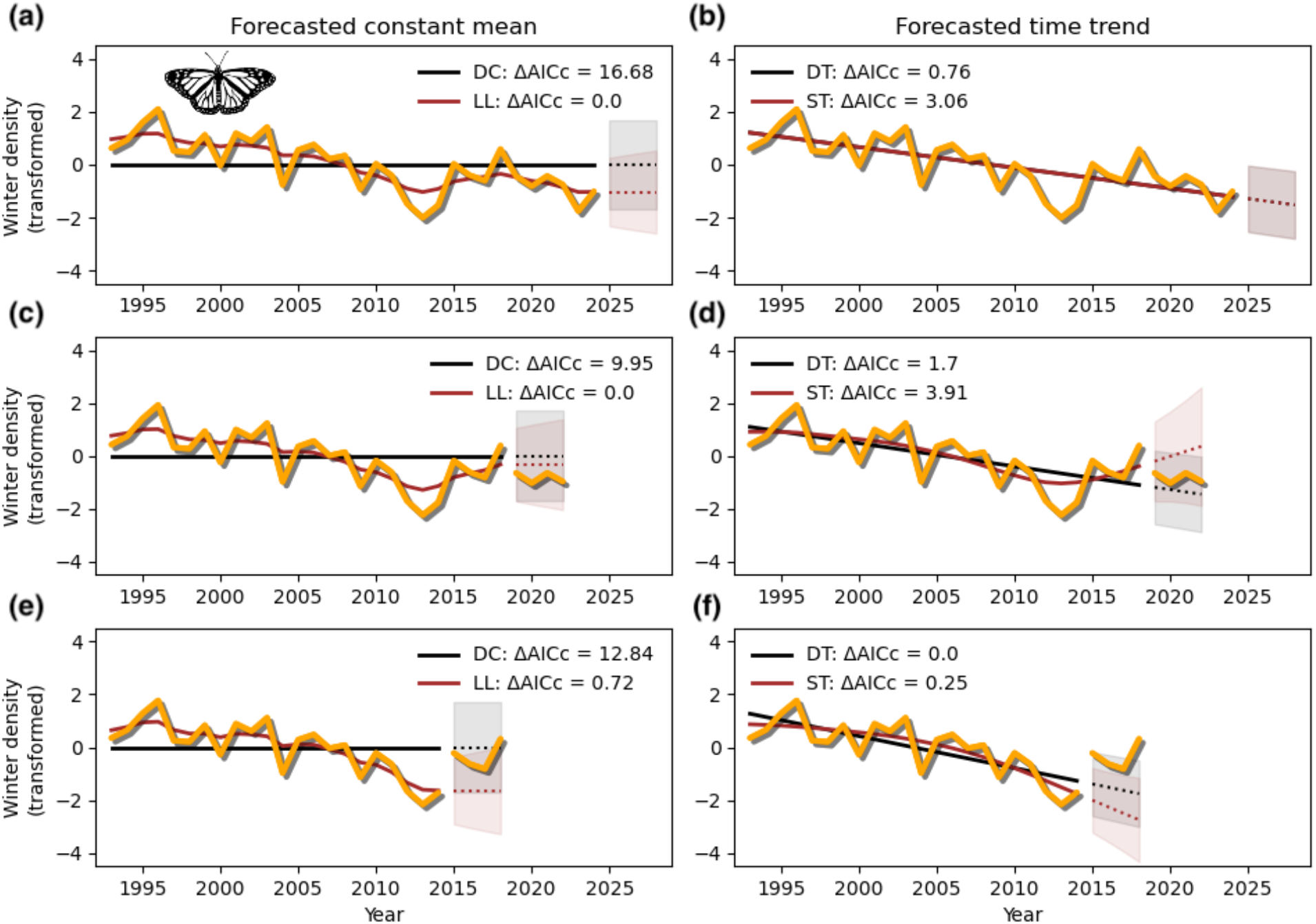
Eastern monarch butterfly population: dynamics and forecasts. Panels (a-b) show the complete time series (up to 2024), along with reconstructed mean trajectories and forecasts based on our four models. Panels (c-d) and (e-f) show the analogous results, however, assuming a shorter timespan (up to 2018 and 2014, respectively). The left column shows the fits of the two models exhibiting a constant forecasted mean: deterministic constant model (DC) and local level model (LL). The right column shows the fits of the two models exhibiting a forecasted linear time trend: deterministic trend model (DT) and smooth trend model (ST). All legends show model name abbreviations along with ΔAICc values, where a value of zero designates the best-fitting model. All forecasts are shown for 4 years ahead, including 90% confidence intervals. Icon: public domain (www.phylopic.org).

Second, AICc as a selection metric covers all aspects of the fit, and it might not always confirm our “visual intuition”. The smooth trend model seems to confirm that after the monarch population hit its bottom in 2013/14, it then increased again (Figure 1d). However, the local level model is the better-fitting model to explain the data up to 2018, and thus it delivers the most parsimonious forecast. In fact (and unfortunately), the increasing period ended in 2018, so the visually intuitive forecast of the smooth trend model was wrong. Third, forecasting is not foreseeing: when only analyzing data up to year 2014, the best-fitting model is the deterministic trend model with a forecasted mean continuing to decline. Nobody could have anticipated the sharp increase in 2015. Here, even if the local level model (as second-best model) suggests a forecasted constant mean, its forecasted variance equally well (or better) captures the unforeseeable turn in dynamics. In sum, selecting the best-fitting model can lead to a forecasted constant mean, despite stochastic increases or decreases in the data.

### Analyzing the high prevalence of forecasted constant means

To investigate what best explains the proportion of time series with forecasted constant mean vs. time trend (see the Results section), we used a random forest classifier [40,41]. An advantage over similar methods like a logistic regression is that the algorithm automatically considers nonlinear and interaction effects among predictors (henceforth called features). We analyzed population-level and community-level results separately. For each set, we split all analyzed time series into a training and a test set, the latter one used for out-of-sample predictions. Because the target data were already unbalanced, we refrained from splitting the time-trend instances into declining vs. increasing categories – doing so would have further skewed the class balance and reduced sample support in each group.

Our selected features cover different aspects of the populations, communities, and the analyzed data. To roughly summarize the time series dynamics, we first computed first-differences (rates of change, RoC) of the transformed time series; from eq. 1a, these are the differences *y*_*t*_ *™ y*_*t ™*1_. Based on these differences, we then computed the mean(RoC) and the var(RoC), and included these two metrics as features. Note that these metrics are model-free summaries computed from the observed data, prior to any UCM fitting. The idea behind these data-related features is that together they help roughly distinguish between dynamics. For example, consider a time series with a linear trend and stochastic deviations around this trend, and a second time series with no time trend but the same stochastic deviations around the constant mean. This will result in var(RoC) being identical for the two time series, but mean(RoC) indicating a difference in the mean dynamics. For nonlinear time trends, likewise both features together roughly summarize the time series dynamics. In sum, we expect mean(RoC) and var(RoC) as system-dynamics metrics to help explain the type of forecasted mean (ESM: Figure S5). To capture “data quality”, we included three features: time series length (henceforth ‘length’), proportion of non-missing values (henceforth ‘p(data)’), and proportion of unique (non-missing) values (henceforth ‘p(unique)’). Finally, to reflect biological signal we included taxonomic information and locality (coordinates). We matched all taxonomic data with the GBIF backbone taxonomy (https://doi.org/10.15468/39omei), and we then used the first four components from a PCA on one-hot-encoded taxonomic data. In ESM: Section S2 we provide more details on data preprocessing and feature selection.

We used the trained random forest classifier to investigate feature importance in terms of predicting a constant vs. linearly changing forecasted mean. Specifically, we computed SHAP values (Shapley Additive ExPlanations values), a widespread approach to explain machine learning models [42,43]. For each instance (e.g. a population) the respective prediction is compared to the overall average prediction, and based on game-theoretic methods the influence of each feature is computed to best explain the difference between the instance-specific prediction and overall average prediction. Advantages over alternative ways to compute feature importance include the additive nature of SHAP values (for example, to group features), their consistency (features that add more value to a prediction yield higher SHAP contributions), and their local reliability (providing instance-level insights that aggregate to global interpretations) [43]. SHAP values have the same units as the target variables: for a classification, they are measured in log-odds. In our analysis, the sign and magnitude of the SHAP values jointly influence the predictions: positive values push predictions towards a forecasted constant mean, and negative values towards a forecasted time trend.

### Importance of biological processes in trend classification

To understand the extent to which biological processes influence trend classification, we followed four lines of evidence. First, we inspected the relative importance of taxonomy and location in predicting trend classification (preceding paragraph). Second, we expected short-term fluctuations to be associated with life history traits and locality ([44], and references therein). Thus, based on available auxiliary data, we refit the random forest classifier for amniote populations (birds, mammals, and reptiles; *n* = 6,916), where we substituted taxonomy-related features (see above) with four life history traits. The life-history traits are age at maturity, maximum lifespan, body mass, and yearly reproductive output [45,46]. In ESM: Section S3 we provide more details on the auxiliary data and data preprocessing. Third, from the SHAP analysis var(RoC) emerged as the most important driver of the type of forecasted mean (see the Results section). As a visual assessment, we examined the distribution of var(RoC) across the Tree of life (64 taxonomic classes). Fourth, to assess how the variance components of the fitted UCMs – observation variance and process variance – relate to var(RoC), we fitted a linear regression in log-log space and compared elasticity values (ESM: Section S3).

## Results

### Forecasted dynamics of populations and communities

After fitting the unobserved components models and retaining time series with no missing attributes for the subsequent classification (next subsection), we grouped time series according to the type of forecasted mean of the respective best-fitting model: forecasted time trend (deterministic trend model and smooth trend model) vs. forecasted constant mean (deterministic constant model and local level model). We found that only 23.5% of populations had forecasted time trends (increasing and decreasing ones roughly 12% each), while 76.5% had a forecasted constant mean (Table 1). For communities, this figure rises to 84.0%, with a negligible difference between temporal *α*- and *β*-diversity time series. The simple baseline model (deterministic constant model) with a forecasted constant mean was selected as best-fitting model for 51.2% of the population-level and 69.6% of the community-level time series (Table 1).

**Table 1.** Overview of forecasted mean results. The columns give: *n*, number of time series; *p*(constant mean), the percentage of time series with a forecasted constant mean; *p*(time trend), the percentage of time series with a forecasted linear trend; *p*(baseline), the percentage of time series in which the baseline model (deterministic constant model) is the best-fitting model.

| | $n$ | $p(\text{constant mean})$ | $p(\text{time trend})$ | $p(\text{baseline})$ |
| --- | --- | --- | --- | --- |
| Populations | 43,403 | 76.5 % | 23.5 % | 51.2 % |
| Communities |  |  |  |  |
| total | 10,409 | 84.0 % | 16.0 % | 69.6 % |
| α-diversity | 5,098 | 84.6 % | 15.4 % | 64.6 % |
| β-diversity | 5,311 | 83.4 % | 16.6 % | 74.7 % |

### Explaining why forecasted constant means are most common

To investigate what biological and data-related characteristics drive the dominance of a forecasted constant mean, we first fit a random forest classifier with the type of forecasted mean as the dependent variable, followed by a SHAP analysis. The trained random forest classifier achieved good predictive ability, with a weighted average f1-score of 0.82 (ESM: Table S2). The model, however, was better at predicting forecasted constant means (class 1; f1-score: 0.88) than forecasted time trends (class 0; f1-score: 0.65), despite explicitly accounting for class imbalance (76.5% vs. 23.5%; Table 1). Of the actual time series, 28.7% with forecasted time trends were misclassified as having a forecasted constant mean, while 14.9% of the time series classified as having a forecasted time trend were actually constant-mean cases. One reason for misclassification might be the diffuse boundaries of feature values between the two classes (ESM: Figure S5). A second reason might be the limited support for time series with forecasted time trends, due to class imbalance (ESM: Table S1). A third reason might be that the features used here are less suited to explain forecasted time trends (see the Discussion).

The SHAP analysis at the population-level showed that the most important feature driving a predicted forecasted constant mean vs. time trend is the variation in the rate of change in the data, var(RoC), followed by mean(RoC) and time series length (Figure 2). The results at the community-level are very similar (Figure 2). For these importance rankings, we computed the mean absolute SHAP value per feature. A more-detailed exploration of the SHAP value results is given in ESM (Figures S6-S12). Both metrics var(RoC) and mean(RoC) – capturing characteristics of variation in the dynamics in the time-series data – account for approximately 50% of the total importance. Time series with high variation in yearly fluctuations are more likely to be associated with a forecasted constant mean, and there is the same association for (very) low-magnitude mean(RoC) values and high-magnitude mean(RoC) values (ESM: Figure S7-S8). Short time series tend to be identified as having forecasted means containing time trends, while longer time series have an increased probability of a forecasted constant mean (ESM: Figure S7-S8).

**Figure 2.**
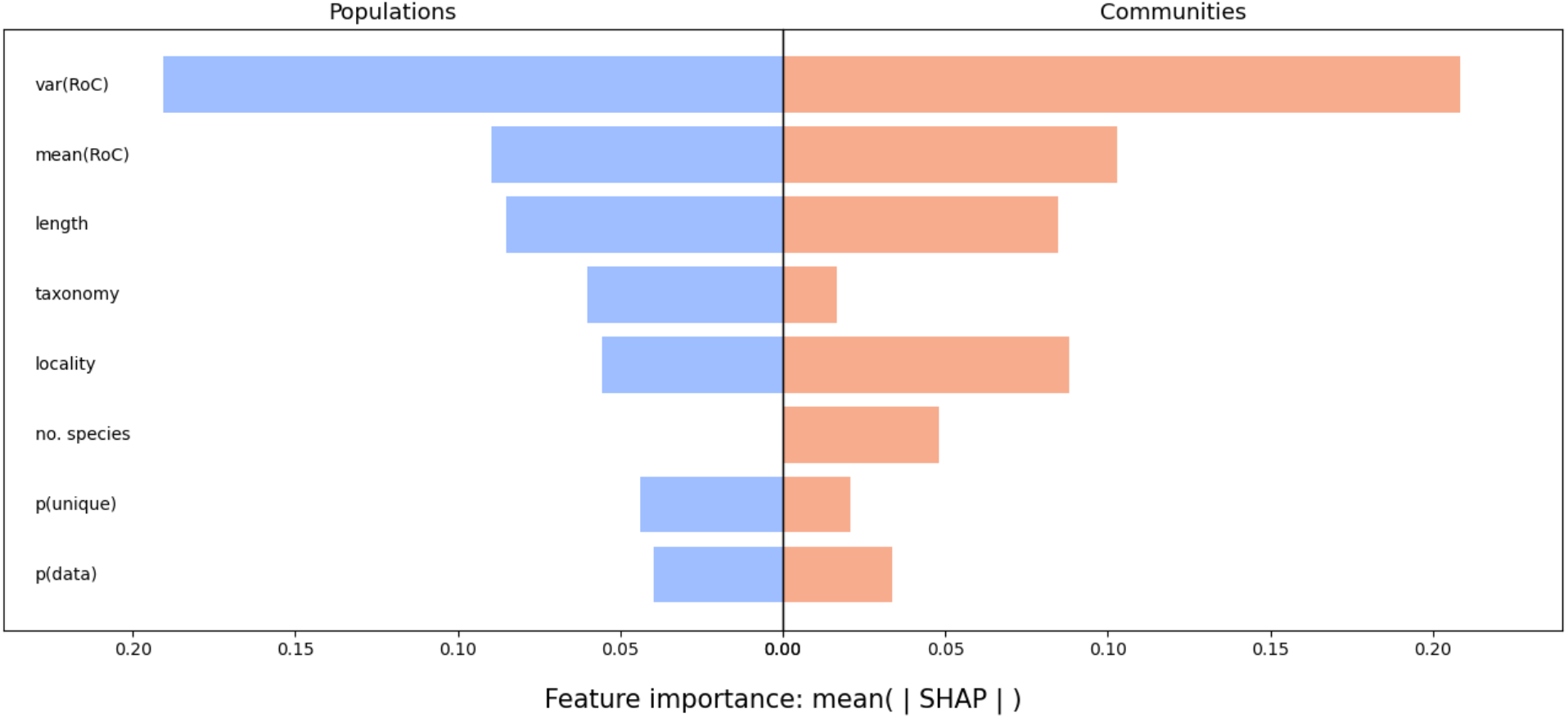
Feature importance values from the random forest classifiers. For each feature in the model, the mean absolute SHAP value is shown separately for population-level and community-level predictions. Note that the features ‘locality’ and ‘taxonomy’ are aggregated (i.e. summed effects): ‘locality’ represents coordinates, and ‘taxonomy’ comprises four PCA components (populations) or one-hot-encoded taxonomic group information (communities).

### Biological structure in trend classification

To assess the extent to which trend classification exhibits biological structure, we followed four lines of evidence. First, taxonomy and location both exceeded the importance of data-quality metrics in the global classifier, retaining meaningful predictive importance after conditioning on var(RoC), mean(RoC), and sampling-related features (Figure 2). This biological structure is visible across the Tree of Life, for example with vertebrates showing a markedly higher proportion of forecasted time trends than invertebrate phyla (Figure 3).

**Figure 3.**
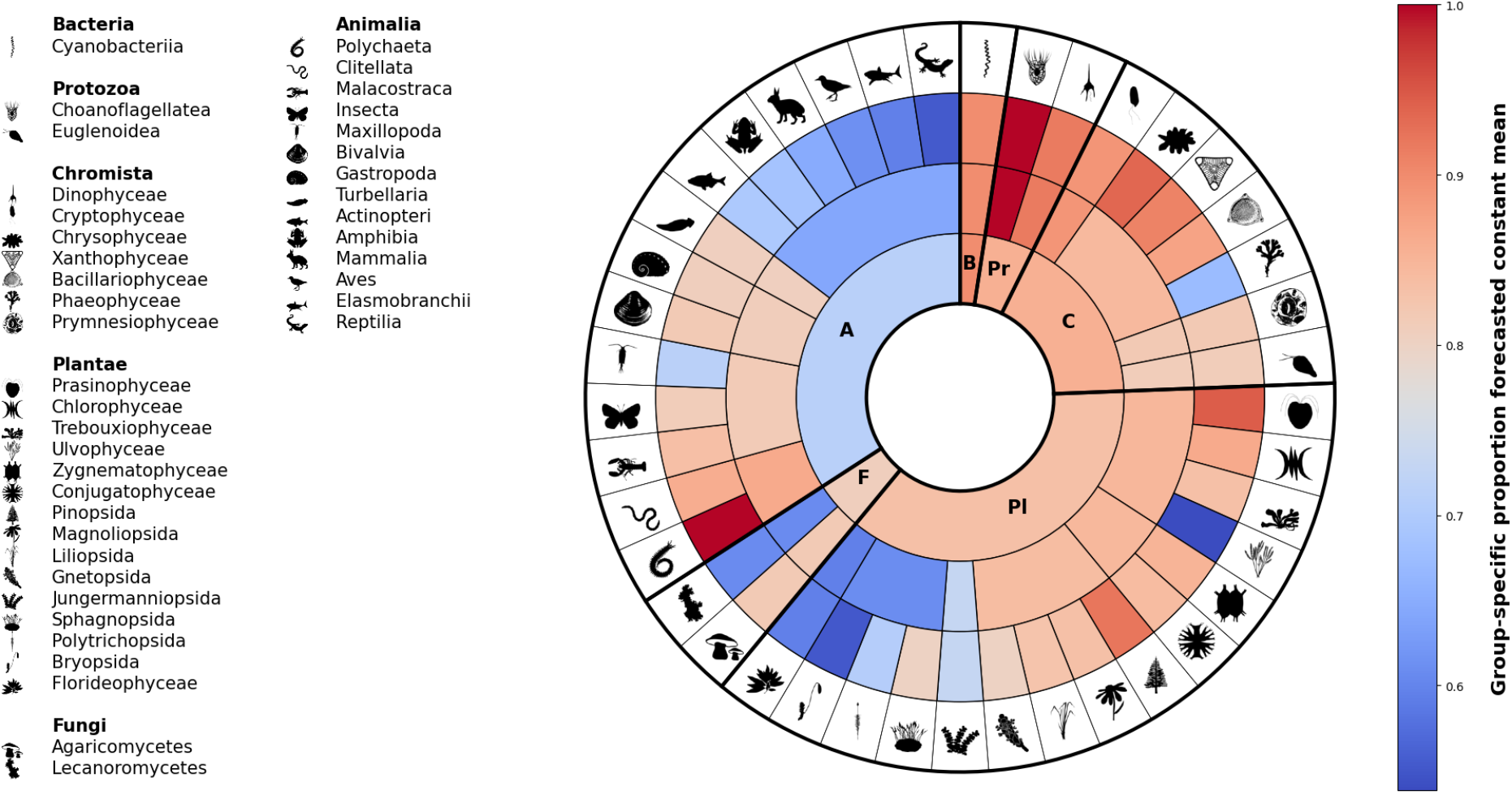
Visualization of the taxonomy dependence of forecasted constant mean. For population-level time series, the nested pie charts show the proportion of forecasted constant means per taxon (ranks kingdom, phylum, class). Only classes with at least ten populations are shown (41 out of 64 classes: ESM: Figure S3). Each ring stepping towards the center shows a higher taxonomic level. All taxa are ordered clockwise in descending order (classes and phyla within higher taxon). Kingdom abbreviations: F[ungi], C[hromista], B[acteria], Pr[otozoa], A[nimalia], Pl[antae]. Icons: public domain (www.phylopic.org).

Second, within amniote populations alone (*n* = 6,916), the classifier achieved a weighted average F1-score of 0.82 (Table S3), with var(RoC) again the dominant predictor. However, life-history traits and location collectively outweighed data-quality metrics in predictive importance (ESM: Figure S13), consistent with the global result but now grounded in life-history traits rather than taxonomy.

Third, a multitude of factors can drive variation in yearly fluctuations, ranging from life-history characteristics to climatic conditions, harvesting, land-use change, and community-specific dynamics. Consistent with this diversity of drivers, var(RoC) itself exhibited pronounced taxonomic structure across the Tree of Life (ESM: Figure S9).

Fourth, var(RoC) scaled more strongly with process variance than observation variance (elasticity ratio = 3.73; RZ = 0.48). This suggests that short-term variability reflects intrinsic ecological dynamics rather than solely observation noise (ESM: Table S4).

## Discussion

Using unobserved components models designed to forecast near-term population and community trends, we investigated the forecasts for 53,812 population- and community-level time series across the globe and the Tree of Life (Table 1). We then investigated what biological and data-related characteristics of the time series best explain the high prevalence of forecasted constant means.

In contrast to the prevailing narrative of pervasive biodiversity decline, our results show that approximately 80% of population and community-level time series have a forecast of no change (Table 1); when a time trend is forecast, increasing trends are as common as decreasing trends, mirroring findings from retrospective analyses ([7], and references therein). Furthermore, the deterministic constant model that assumes fluctuations around a constant mean (Figure 1) was selected as best-fitting model for ∼50% of the population and ∼70% of the community-level time series (Table 1). This result echoes previous findings from forecasts for bird species [19]. As illustrated with the Eastern monarch butterfly population (Figure 1), we emphasize that the visual appearance of a trend in the data is often not upheld by a statistical model that correctly accounts for variation and consequently forecasts no future change in the mean.

From a methodological perspective, unobserved components models in state-space form are robust statistical tools with high statistical power to identify trends (e.g. [36]), and therefore our results are not the consequence of inadequate methods. Moreover, random forest models in conjunction with SHAP analysis give a sensible and robust approach to investigate complex data such as the drivers of patterns identified by the unobserved components models (e.g. [43]). Nonetheless, our time-series analyses have not addressed spatial and taxonomic correlation among the time series [8], which would become important when computing an overall expected trend for a specific taxonomic group (e.g. [47]). In general, accounting for spatial and taxonomic correlation will decrease the likelihood of identifying significant trends [48], leading to the categorization of even more forecasted constant means. Further, we analyzed each time series without including covariates, because it would be impractical to try to assemble appropriate covariate time series for the large number of time series we analyzed. While including covariates in the time-series analyses may have revealed interesting patterns, we do not think that this would change our finding of few time series with forecasted time trends, because the simplest model of a constant mean was most often the best-fitting model (see also [17]).

We also addressed the importance of time series length. Many studies call for longer time series (e.g. [49,50]). Our results at first sight seem to challenge this call: time series shorter than 15 years were more likely associated with forecasted time trends (ESM: Figure S6-S7). However, this association is mainly due to spurious trends in the short time series. Beyond 15 years, the likelihood of a forecasted constant mean increases, and mean(RoC) often approaches zero. Longer time series are always preferable, both for narrowing the uncertainty in mean(RoC) and var(RoC), and for ensuring robust analyses. However, this will often result in a forecasted constant mean for the best-fitting models. This observation mirrors retrospective trend analyses, which indicate that steady means are common ([7], and references therein). We should be wary of short time series because their forecasted trends more often than not may be spurious.

What do our findings imply for near-term biodiversity forecasts? First, low variation in the rate of change, var(RoC) – the strongest predictor of a forecasted time trend (Figure 2) – is the exception, not the rule in ecological data (ESM: Figure S7-S8), so forecasts of no change should be common. Second, the distribution of data attributes in current ecological data collections – time-series length, sampling unit and frequency, and others – is unlikely to change soon. These constraints will continue to hinder our ability to confidently detect and forecast true underlying time trends. Third, biological structure (taxonomy, location, and life-history traits) consistently predicted trend classification beyond var(RoC) and data-quality metrics (Figures 2-3, ESM: Fig. S13). Further, var(RoC) was more strongly associated with process variance than observation variance, suggesting it captures intrinsic ecological dynamics more than observation noise (ESM: Table S4). Moreover, some taxonomic groups (e.g., vertebrates) tend to have lower var(RoC), so taxon-specific forecasting can be more informative when the forecasting scope is limited (ESM: Figure S9). Obviously, vertebrates represent only one phylum among many in the Tree of Life, and relying on their trends is misleading for taxonomic generalizations. When managing or restoring whole ecosystems, the entire taxonomic breadth should be used to produce forecasts of expected changes in composition and function. Even so, population-level – and especially community-level – analyses across the Tree of Life will often yield a forecasted constant mean (Table 1). Finally, our result that roughly 80% of populations and communities have forecasts of no change is consistent with retrospective work ([7], and references therein), suggesting that persistent directional trends may be less widespread across the Tree of Life than commonly assumed.

The ongoing biodiversity crisis demands both retrospective and prospective trend analyses. Working with typical ecological time-series data can be challenging, but overall a nuanced picture is emerging of the extent of population- and community-level changes. Near-term statistical forecasts are critical to inform stakeholders, as well as devising and monitoring conservation actions. Despite their simplicity, we believe parsimonious forecasting approaches continue to offer valuable, robust, and actionable insights, to address the challenges posed by a changing natural world.

## Supporting information

ESM

## Acknowledgments

We thank Simon Egli for sharing his long-term macrofungal data.

## Data availability

Most of the data that support the findings of this study are publicly available online and are referenced in the main text. See Appendix S1: Table S1 for further details.

### Code availability

Code (Python), along with raw results files, will be uploaded to the zenodo repository after acceptance.

### Author contributions

**C.B**. Conceptualization, Methodology, Formal analysis, Data Curation,

Writing - Original Draft, Writing - Review & Editing, Visualization. **A.R.I**. Methodology, Formal analysis, Writing -Review & Editing.

## References

1. Wilson EO. 1992 The Diversity of Life. Harvard University Press.

2. Dirzo R, Young HS, Galetti M, Ceballos G, Isaac NJB, Collen B. 2014 Defaunation in the Anthropocene. Science 345, 401–406. (doi:10.1126/science.1251817)

3. Wiens JJ, Saban KE. 2025 Questioning the sixth mass extinction. Trends in Ecology & Evolution 40, 375–384. (doi:10.1016/j.tree.2025.01.002)

4. Rosenberg KV et al. 2019 Decline of the North American avifauna. Science 366, 120–124. (doi:10.1126/science.aaw1313)

5. UNEP-WCMC. 2024 State of the World’s Migratory Species Report.

6. Hallmann CA et al. 2017 More than 75 percent decline over 27 years in total flying insect biomass in protected areas. PLOS ONE 12, e0185809. (doi:10.1371/journal.pone.0185809)

7. Dornelas M et al. 2023 Looking back on biodiversity change: lessons for the road ahead. Philosophical Transactions of the Royal Society B: Biological Sciences 378, 20220199. (doi:10.1098/rstb.2022.0199)

8. Johnson TF et al. 2024 Revealing uncertainty in the status of biodiversity change. Nature, 1–7. (doi:10.1038/s41586-024-07236-z)

9. Boënnec M, Dakos V, Devictor V. 2024 Sources of confusion in global biodiversity trends. Oikos 2024, e10732. (doi:10.1111/oik.10732)

10. Toszogyova A, Smyčka J, Storch D. 2024 Mathematical biases in the calculation of the Living Planet Index lead to overestimation of vertebrate population decline. Nat Commun 15, 5295. (doi:10.1038/s41467-024-49070-x)

11. Urban MC et al. 2016 Improving the forecast for biodiversity under climate change. Science 353, aad8466. (doi:10.1126/science.aad8466)

12. Dietze M et al. 2024 Near-term ecological forecasting for climate change action. Nat. Clim. Chang. 14, 1236–1244. (doi:10.1038/s41558-024-02182-0)

13. Newbold T et al. 2015 Global effects of land use on local terrestrial biodiversity. Nature 520, 45–50. (doi:10.1038/nature14324)

14. IPBES. 2019 Global assessment report on biodiversity and ecosystem services of the Intergovernmental Science-Policy Platform on Biodiversity and Ecosystem Services. (doi:10.5281/zenodo.6417333)

15. Wood KA, Stillman RA, Hilton GM. 2018 Conservation in a changing world needs predictive models. Animal Conservation 21, 87–88. (doi:10.1111/acv.12371)

16. Tulloch AIT, Hagger V, Greenville AC. 2020 Ecological forecasts to inform near-term management of threats to biodiversity. Global Change Biology 26, 5816–5828. (doi:10.1111/gcb.15272)

17. Lasky JR, Hooten MB, Adler PB. 2020 What processes must we understand to forecast regional-scale population dynamics? Proceedings of the Royal Society B: Biological Sciences 287, 20202219. (doi:10.1098/rspb.2020.2219)

18. Lewis ASL et al. 2022 Increased adoption of best practices in ecological forecasting enables comparisons of forecastability. Ecological Applications 32, e2500. (doi:10.1002/eap.2500)

19. Harris DJ, Taylor SD, White EP. 2018 Forecasting biodiversity in breeding birds using best practices. PeerJ 6, e4278. (doi:10.7717/peerj.4278)

20. Olsson F et al. 2025 What can we learn from 100,000 freshwater forecasts? A synthesis from the NEON Ecological Forecasting Challenge. Ecological Applications 35, e70004. (doi:10.1002/eap.70004)

21. Ward EJ, Holmes EE, Thorson JT, Collen B. 2014 Complexity is costly: a meta-analysis of parametric and non-parametric methods for short-term population forecasting. Oikos 123, 652–661. (doi:10.1111/j.1600-0706.2014.00916.x)

22. Pennekamp F et al. 2019 The intrinsic predictability of ecological time series and its potential to guide forecasting. Ecological Monographs 89, e01359. (doi:10.1002/ecm.1359)

23. Bozzuto C, Ives AR. 2024 Predictability of ecological and evolutionary dynamics in a changing world. Proceedings of the Royal Society B: Biological Sciences 291, 20240980. (doi:10.1098/rspb.2024.0980)

24. Dornelas M et al. 2025 BioTIME 2.0: Expanding and Improving a Database of Biodiversity Time Series. Global Ecology and Biogeography 34, e70003. (doi:10.1111/geb.70003)

25. Filippova N, Rudykina E, Zvyagina E, Dobrynina A, Lutovinova V, Popesku K. 2024 Plot-based observations of terrestrial macrofungi in different forest types of boreal zone in West Siberia (2015-2024). (doi:10.15468/ge1hkl)

26. Prendergast J, Bazeley-White E, Smith O, Lawton J, Inchausti P, Kidd D, Knight S. 2010 The Global Population Dynamics Database. (doi:10.5063/F1BZ63Z8)

27. Loh J, Green RE, Ricketts T, Lamoreux J, Jenkins M, Kapos V, Randers J. 2005 The Living Planet Index: using species population time series to track trends in biodiversity. Philosophical Transactions of the Royal Society B: Biological Sciences 360, 289–295. (doi:10.1098/rstb.2004.1584)

28. Richter D. 2023 Mycorrhizal Fungi of Native Red Pine Stands in the Forests of the Huron Mountains (1996-2015). (doi:10.6073/pasta/37cf0ded54b059cd46eb25ee529af511)

29. Hacket-Pain A et al. 2022 MASTREE+: Time-series of plant reproductive effort from six continents. Global Change Biology 28, 3066–3082. (doi:10.1111/gcb.16130)

30. Comte L et al. 2021 RivFishTIME: A global database of fish time-series to study global change ecology in riverine systems. Global Ecology and Biogeography 30, 38–50. (doi:10.1111/geb.13210)

31. Welti EAR et al. 2024 Time series of freshwater macroinvertebrate abundances and site characteristics of European streams and rivers. Sci Data 11, 601. (doi:10.1038/s41597-024-03445-3)

32. Pomati F, Shurin JB, Andersen KH, Tellenbach C, Barton AD. 2020 Interacting Temperature, Nutrients and Zooplankton Grazing Control Phytoplankton Size-Abundance Relationships in Eight Swiss Lakes. Front. Microbiol. 10. (doi:10.3389/fmicb.2019.03155)

33. Egli S. 2009 Mykorrhizapilze auf dem Rückzug – was bedeutet das für den Wald? Langzeitforschung für eine nachhaltige Waldnutzung, 51–58.

34. Dornelas M, Gotelli NJ, McGill B, Shimadzu H, Moyes F, Sievers C, Magurran AE. 2014 Assemblage Time Series Reveal Biodiversity Change but Not Systematic Loss. Science 344, 296–299. (doi:10.1126/science.1248484)

35. Cha S-H. 2007 Comprehensive Survey on Distance/Similarity Measures Between Probability Density Functions. Int. J. Math. Model. Meth. Appl. Sci. 1.

36. Harvey AC, Koopman SJ, Penzer J. 1997 Messy Time Series: A Unified Approach. STICERD-Econometrics Paper Series

37. Durbin T late J, Koopman SJ. 2012 Time Series Analysis by State Space Methods. Second Edition. Oxford, New York: Oxford University Press.

38. Auger-Méthé M et al. 2021 A guide to state–space modeling of ecological time series. Ecological Monographs 91, e01470. (doi:10.1002/ecm.1470)

39. Zylstra ER, Ries L, Neupane N, Saunders SP, Ramírez MI, Rendón-Salinas E, Oberhauser KS, Farr MT, Zipkin EF. 2021 Changes in climate drive recent monarch butterfly dynamics. Nat Ecol Evol 5, 1441–1452. (doi:10.1038/s41559-021-01504-1)

40. Breiman L. 2001 Random Forests. Machine Learning 45, 5–32. (doi:10.1023/A:1010933404324)

41. Cutler DR, Edwards Jr. TC, Beard KH, Cutler A, Hess KT, Gibson J, Lawler JJ. 2007 Random Forests for Classification in Ecology. Ecology 88, 2783–2792. (doi:10.1890/07-0539.1)

42. Lundberg SM et al. 2020 From local explanations to global understanding with explainable AI for trees. Nat Mach Intell 2, 56–67. (doi:10.1038/s42256-019-0138-9)

43. Molnar C. 2025 Interpretable Machine Learning: A Guide for Making Black Box Models Explainable. 3rd edn. See https://christophm.github.io/interpretable-ml-book.

44. Albaladejo-Robles G, Böhm M, Newbold T. 2023 Species life-history strategies affect population responses to temperature and land-cover changes. Global Change Biology 29, 97–109. (doi:10.1111/gcb.16454)

45. Myhrvold NP, Baldridge E, Chan B, Sivam D, Freeman DL, Ernest SKM. 2015 An amniote life-history database to perform comparative analyses with birds, mammals, and reptiles. Ecology 96, 3109–3109. (doi:10.1890/15-0846R.1)

46. Pacifici M, Santini L, Marco MD, Baisero D, Francucci L, Marasini GG, Visconti P, Rondinini C. 2013 Generation length for mammals. Nature Conservation 5, 89–94. (doi:10.3897/natureconservation.5.5734)

47. WWF. 2024 Living Planet Report 2024-A System in Peril.

48. Ives AR. 2022 Random errors are neither: On the interpretation of correlated data. Methods in Ecology and Evolution 13, 2092–2105. (doi:10.1111/2041-210X.13971)

49. White ER. 2019 Minimum Time Required to Detect Population Trends: The Need for Long-Term Monitoring Programs. BioScience 69, 40–46. (doi:10.1093/biosci/biy144)

50. Cusser S, Helms IV J, Bahlai CA, Haddad NM. 2021 How long do population level field experiments need to be? Utilising data from the 40-year-old LTER network. Ecology Letters 24, 1103–1111. (doi:10.1111/ele.13710)

