## Supplementary material for "Forecasts for 80% of wild populations and communities predict no change, and why": ESM

### Electronic supplementary material

This PDF-file contains:

- Supplementary Tables S1 – S4
- Supplementary Figures S1 – S12
- Supplementary Methods: Sections S1 – S3

#### Supplementary Tables

**Table S1**. Data collections and number of time series used in this study. Data collections: BioTIME[(1)]; GBIF[(2)]; GPDD[(3)]; LPI[(4)]; LTER[(5)]; MASTREE+[(6)]; TREAM[(8)]; RivFISH[(9)]; Phytoplankton[(9)]; Swiss fungi[(10)]. Abbreviations: C = community-level time series; P = population-level time series.

| | | BioTIME | GBIF | GPDD | LPI | LTER | MASTREE+ | TREAM | RivFISH | Phyto-plankton | Swiss fungi |
|---|---|---|---|---|---|---|---|---|---|---|---|
| Amphibians | C | | | | | | | | | | |
| | P | | | | 255 | | | | | | |
| Birds | C | 1,440 | | | | | | | | | |
| | P | | | | 7,372 | | | | | | |
| Fish | C | | | | | | | | 4,106 | | |
| | P | | | | 5,055 | | | | | | |
| Mammals | C | 1,807 | | | | | | | | | |
| | P | | | | 1,802 | | | | | | |
| Reptiles | C | | | | | | | | | | |
| | P | | | | 378 | | | | | | |
| Invertebrates, freshwater | C | | | | | | | 872 | | | |
| | P | | | | | | | 8,662 | | | |
| Invertebrates, terrestrial | C | 1,009 | | | | | | | | | |
| | P | | | 1,412 | | | | | | | |
| Plants | C | 1,175 | | | | | | | | | |
| | P | 13,256 | | | | | 3,302 | | | | |
| Phytoplankton | C | | | | | | | | | | |
| | P | | | | | | | | | 1,310 | |
| Fungi | C | | | | | | | | | | |
| | P | 40 | 49 | | | 66 | | | | | 444 |

[(1)] https://biotime.st-andrews.ac.uk/ (accessed 30-10-2024).

[(2)] https://www.gbif.org/dataset/7648036d-fdac-4ebe-8861-9a28134adae1 (accessed 30-03-2025).

[(3)] https://knb.ecoinformatics.org/view/doi:10.5063/F1BZ63Z8 (accessed 01-04-2022).

[(4)] https://stats.livingplanetindex.org/data_portal (accessed 17-02-2025).

[(5)] https://portal.edirepository.org/nis/mapbrowse?packageid=edi.1470.1 (accessed 30-03-2025).

[(6)] https://github.com/JJFoest/MASTREEplus (accessed 07-04-2025).

[(7)] https://knb.ecoinformatics.org/view/doi:10.5063/F1NG4P4R (accessed 30-10-2024).

[(8)] https://zenodo.org/records/13825561 (accessed 11-04-2025).

[(9)] https://zenodo.org/records/13825561 (accessed 11-04-2025).

[(10)] Received from Simon Egli (data owner) on 16-04-2025.

**Table S2**. **Classification reports for population-level and community-level random forest classifiers.** For the random forest classifiers fitted to the population-level and community-level datasets separately, the table shows the respective classification report. In brackets we further report results from 5-fold cross-validation (mean ± std).

| | **precision** | **recall** | **f1-score** | **support** |
|---|---|---|---|---|
| *Population-level* | | | | |
| **mean with time trend** | 0.60<br>(0.612 ± 0.008) | 0.71<br>(0.718 ± 0.008) | 0.65<br>(0.66 ± 0.007) | 2044 |
| **constant mean** | 0.91<br>(0.910 ± 0.0) | 0.85<br>(0.858 ± 0.004) | 0.88<br>(0.882 ± 0.004) | 6637 |
| weighted average | 0.83<br>(0.840 ± 0.0) | 0.82<br>(0.826 ± 0.005) | 0.82<br>(0.832 ± 0.004) | 8681 |
| *Community-level* | | | | |
| **mean with time trend** | 0.44<br>(0.458 ± 0.012) | 0.65<br>(0.673 ± 0.008) | 0.52<br>(0.545 ± 0.010) | 333 |
| **constant mean** | 0.93<br>(0.932 ± 0.002) | 0.84<br>(0.848 ± 0.007) | 0.88<br>(0.888 ± 0.004) | 1749 |
| weighted average | 0.85<br>(0.856 ± 0.003) | 0.81<br>(0.820 ± 0.006) | 0.83<br>(0.833 ± 0.005) | 2082 |

**Table S3**. **Classification reports for the amniote population-level random forest classifier**. The table shows the classification report for the random forest classifier fitted to amniote populations. In brackets we further report results from 5-fold cross-validation (mean ± std).

| | **precision** | **recall** | **f1-score** | **support** |
|---|---|---|---|---|
| *Population-level* | | | | |
| **mean with time trend** | 0.79<br>(0.763 ± 0.009) | 0.76<br>(0.766 ± 0.013) | 0.78<br>(0.764 ± 0.008) | 565 |
| **constant mean** | 0.84<br>(0.838 ± 0.007) | 0.86<br>(0.835 ± 0.008) | 0.85<br>(0.837 ± 0.005) | 819 |
| weighted average | 0.82<br>(0.807 ± 0.006) | 0.82<br>(0.807 ± 0.006) | 0.82<br>(0.807 ± 0.006) | 1384 |

**Table S4**. **OLS log-log regression of var(RoC) on UCM variance components.** The dependent variable is log(var(RoC) + $\varepsilon$), where $\varepsilon = 10^{-8}$ is a small constant added to avoid log(0). "Signal" means level variance ($\sigma_{\xi}^2$) or slope variance ($\sigma_{\zeta}^2$); see eq. 1a-c. The dummy predictor captures the discrete shift for series where process variance is structurally zero. Standard errors are HC3-robust. Elasticity ratio $\beta$(log(signal)) / $\beta$(log($\sigma_{\varepsilon}^2$)) = 3.73. All $p$-values < 0.001.

| Predictor | Estimate | SE | z | 95% CI |
|---|---|---|---|---|
| Intercept | 0.609 | 0.019 | 32.87 | [0.573, 0.646] |
| log($\sigma_{\varepsilon}^2$) | 0.103 | 0.002 | 56.66 | [0.099, 0.107] |
| log(signal) | 0.385 | 0.008 | 47.31 | [0.369, 0.401] |
| Indicator variable (model type) | 6.889 | 0.134 | 51.48 | [6.627, 7.151] |

#### Supplementary Figures

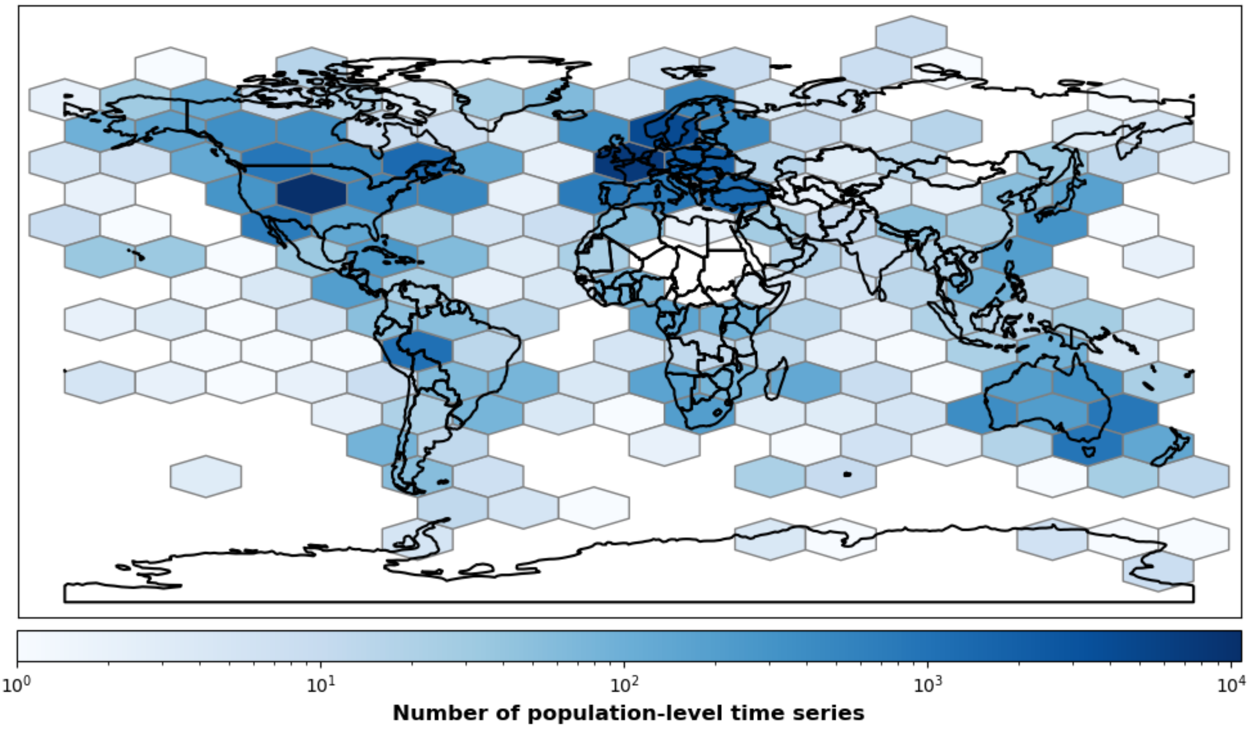


**Figure S1. Geographic distribution of analyzed population-level time series**. The localities of all populations ($n$ = 43,403) are visualized using hexagonal bins, colored according to the number of populations contained (colorbar).

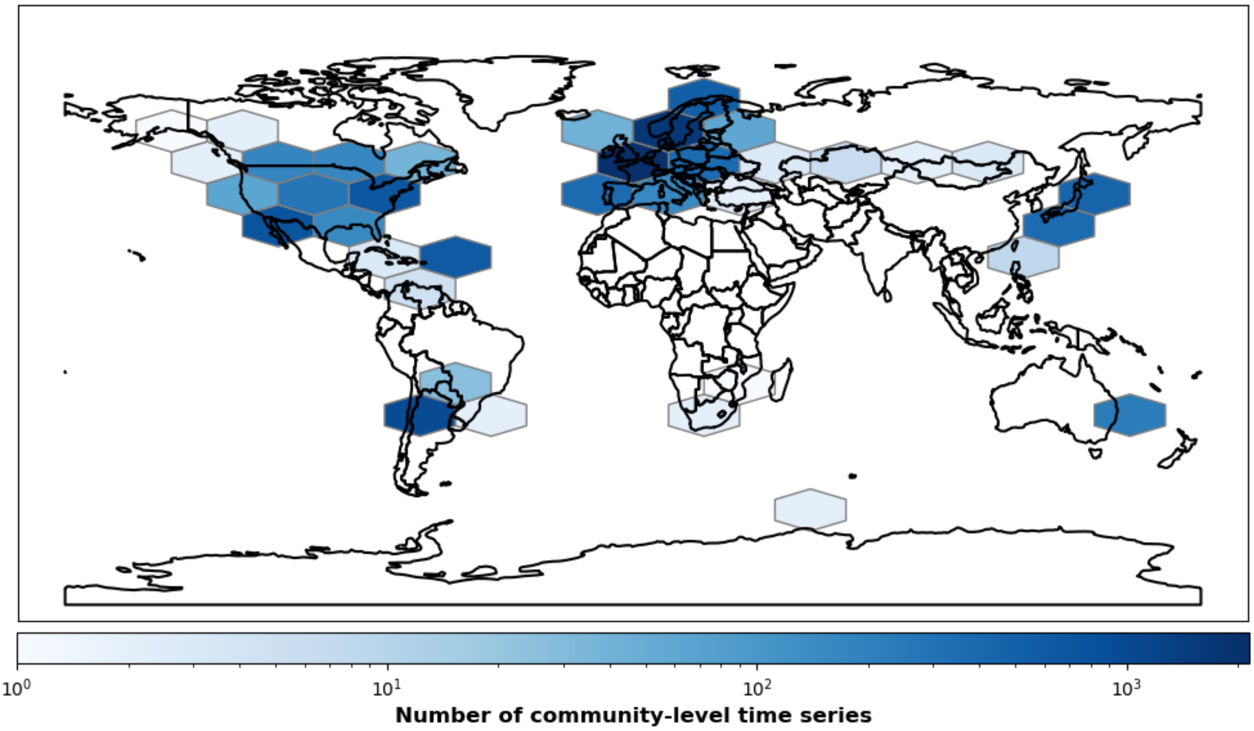


**Figure S2. Geographic distribution of analyzed community-level time series**. The localities of all communities ($n$ = 10,409) are visualized using hexagonal bins, colored according to the number of communities contained (colorbar).

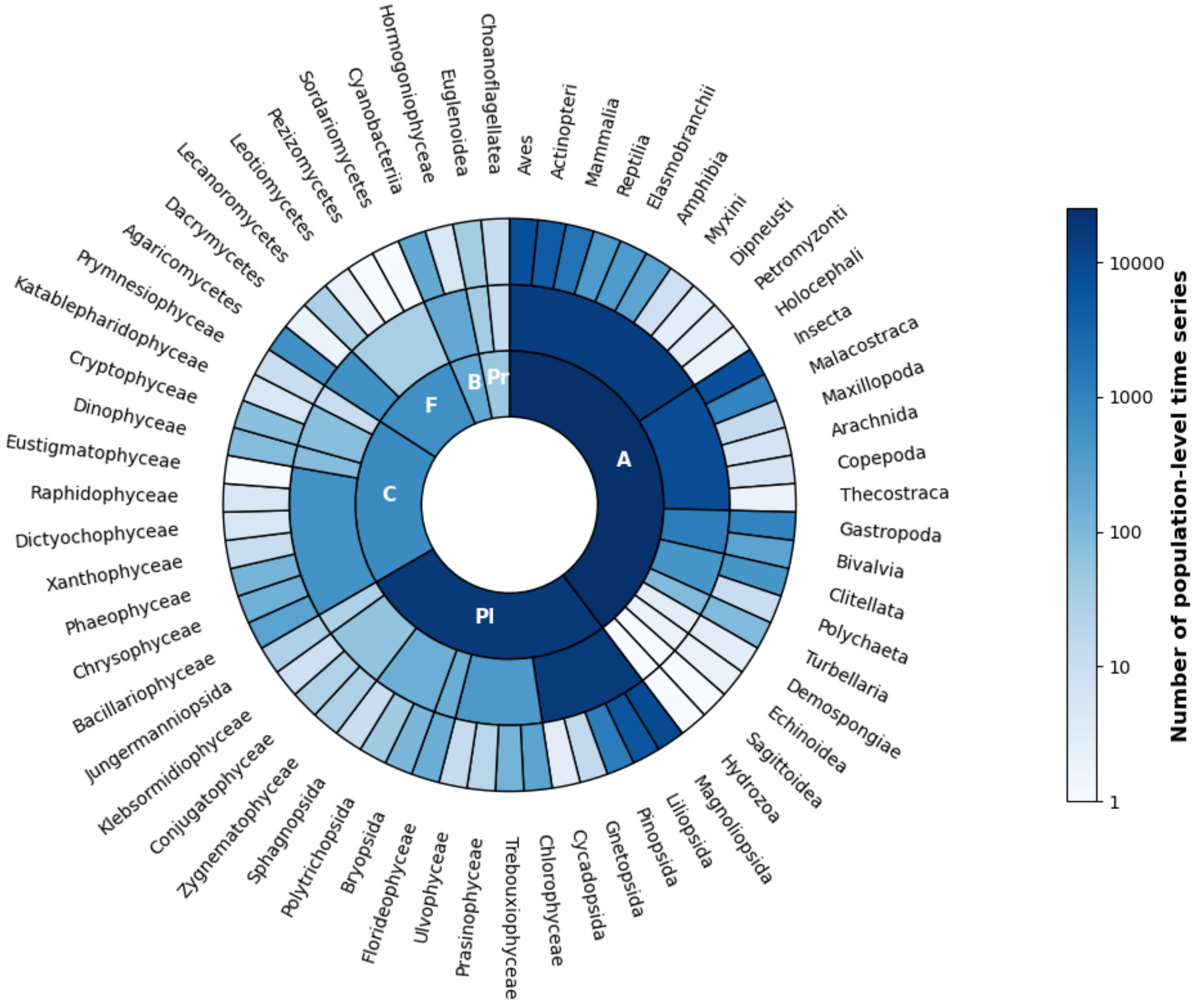


**Figure S3. Taxonomic coverage of analyzed population-level time series**. The nested pie chart shows the number of time series per taxon (colorbar), analyzed at the population level. All taxa are shown in descending order (clockwise; classes and phyla within higher taxon). Kingdom abbreviations: A[nimalia], Pl[antae], C[hromista], F[ungi], B[acteria], Pr[otozoa].

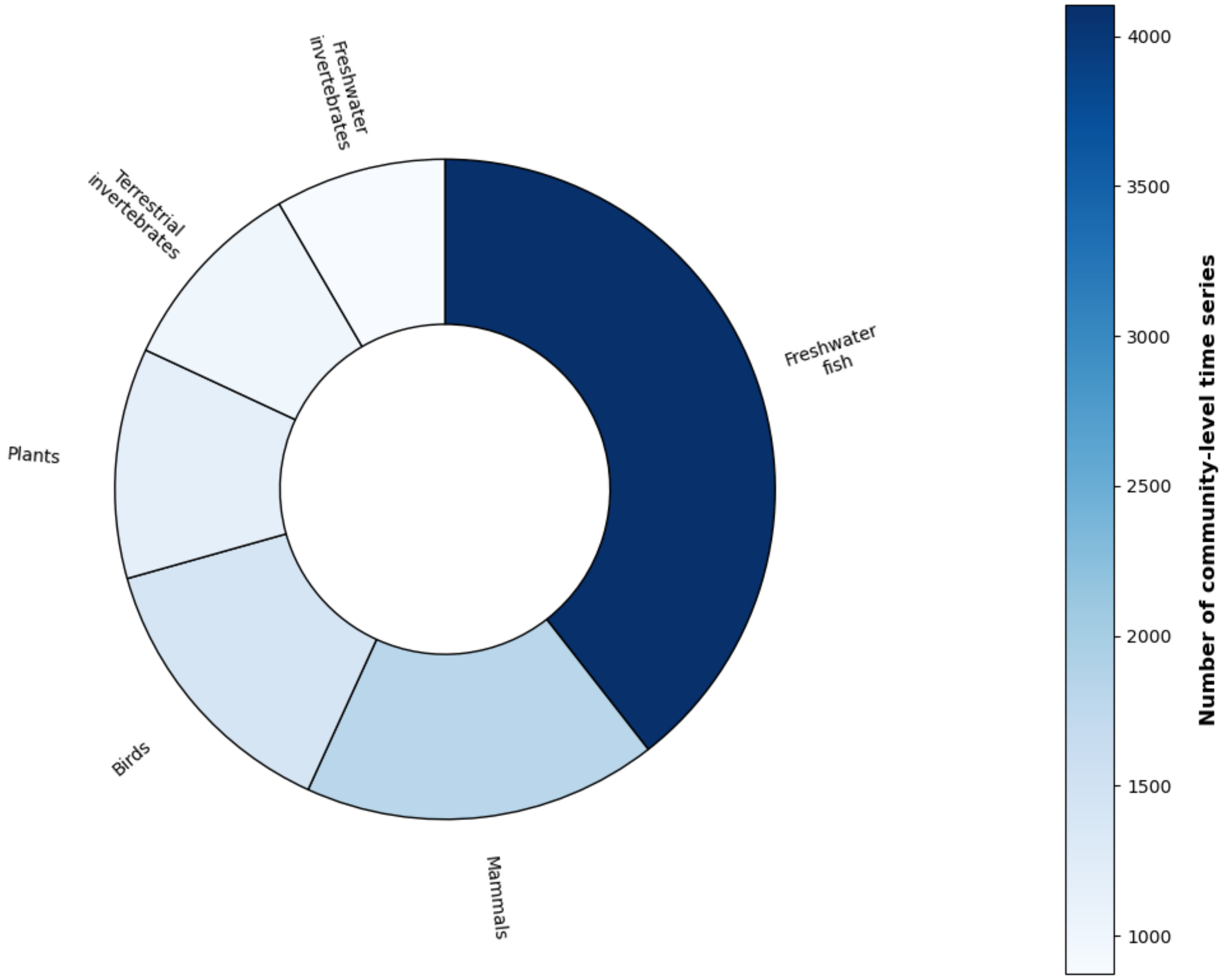


**Figure S4. Taxonomic coverage of analyzed community-level time series**. The pie chart shows the number of time series per taxonomic group (colorbar) at the community level analyzed in this study. All taxonomic groups are ordered in descending order (clockwise).

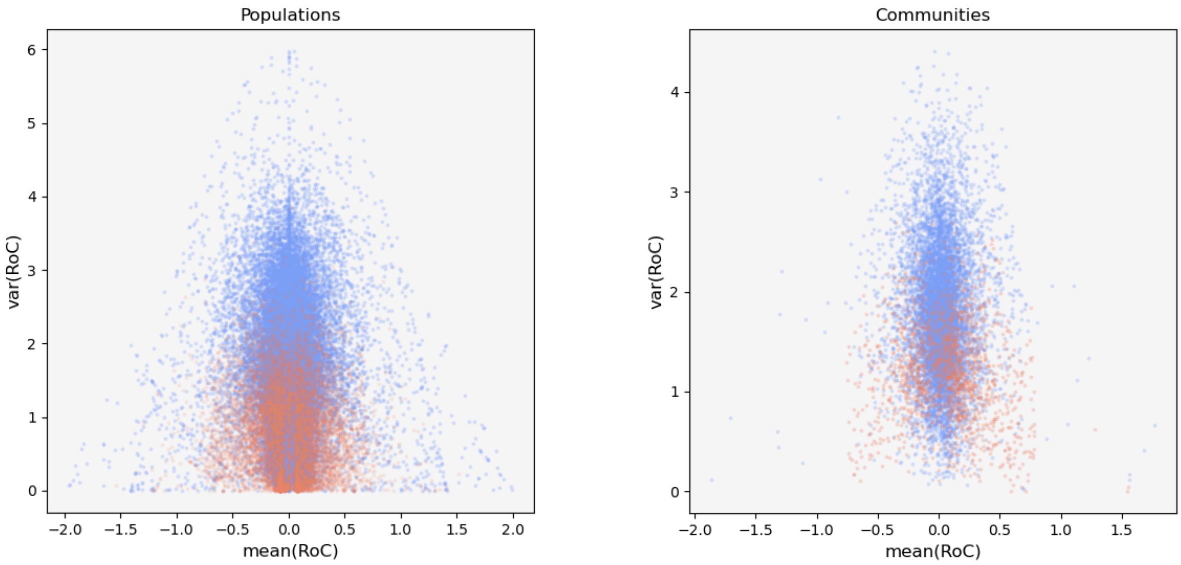


**Figure S5. Empirical relationship between mean(RoC), var(RoC) and type of forecasted mean**. Both panels show the clustering and partial overlap of mean(RoC) and var(RoC) values for all analyzed time series with forecasted constant mean (blue) and forecasted time trend (red), at the population (left) and community level (right).

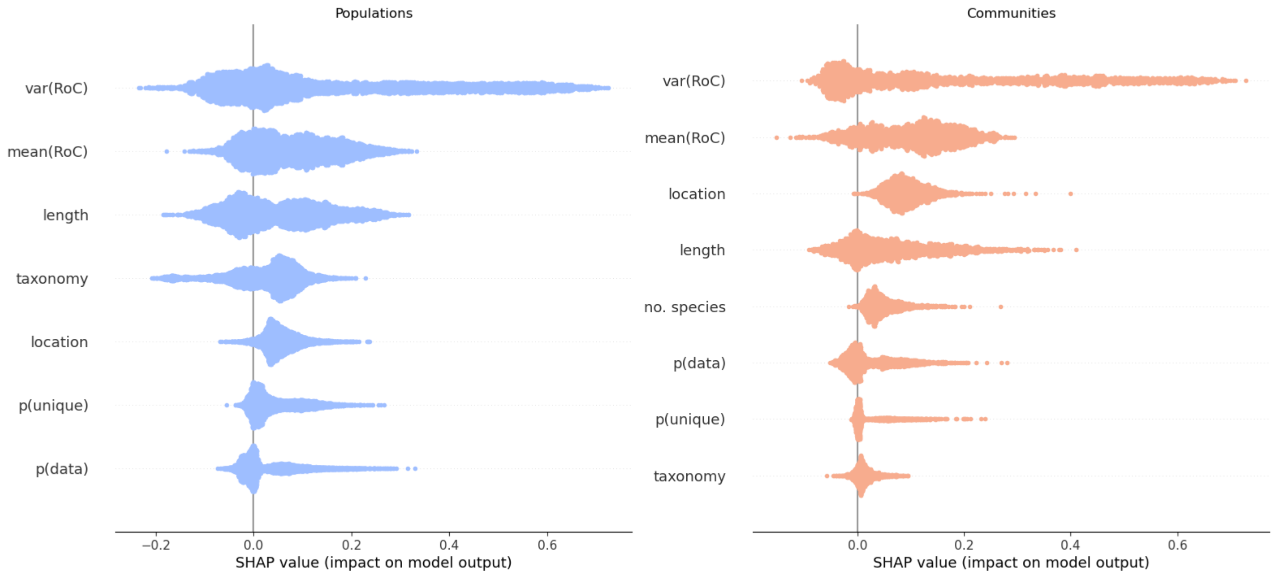


**Figure S6. Summary plots of SHAP values**. Distribution of SHAP values for population-level (left) and community-level random forest classifiers (right). Within each panel, the features are shown in descending order of importance.

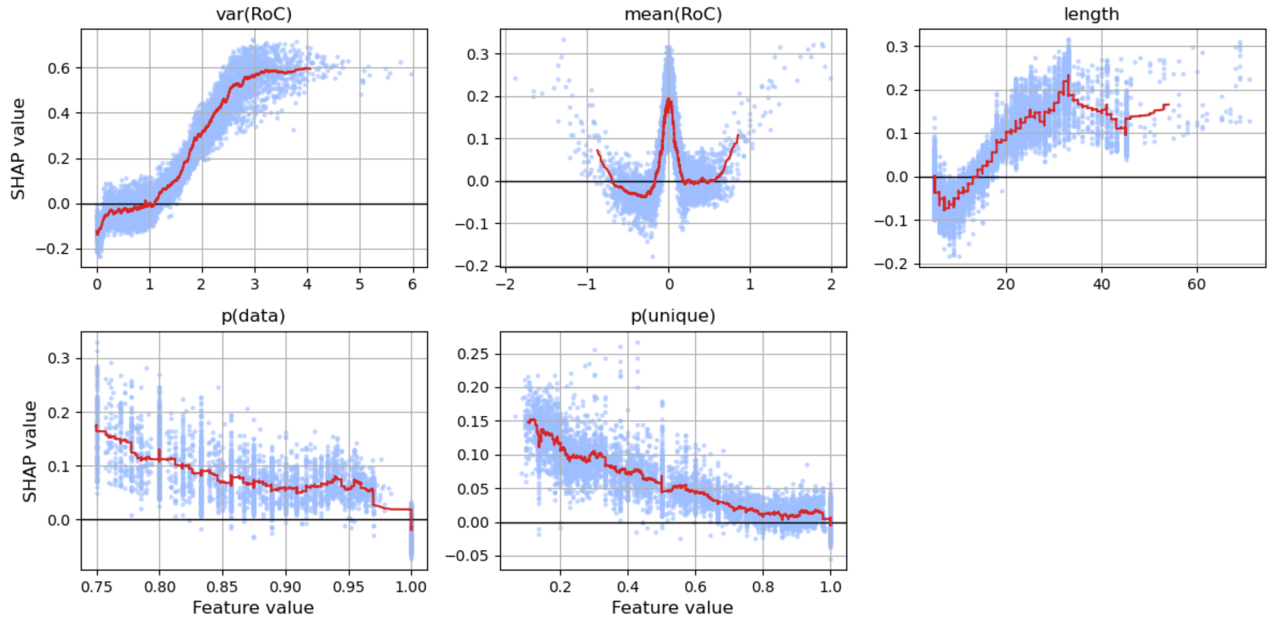


**Figure S7. SHAP values per feature: population-level time series**. For non-aggregated features, each panel shows the distribution of SHAP values as a function of the respective feature values. SHAP values are displayed with dots, and the red curves have been generated with a rolling window.

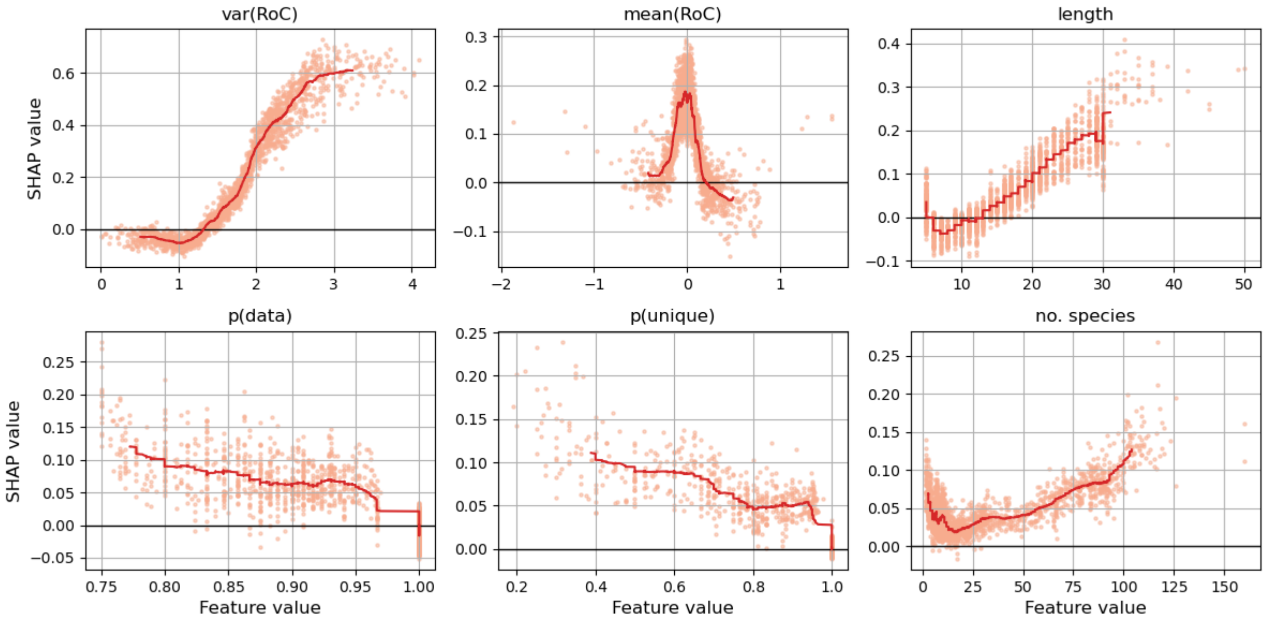


**Figure S8. SHAP values per feature: community-level time series**. For non-aggregated features, each panel shows the distribution of SHAP values as a function of the respective feature values. SHAP values are displayed with dots, and the red curves have been generated with a rolling window.

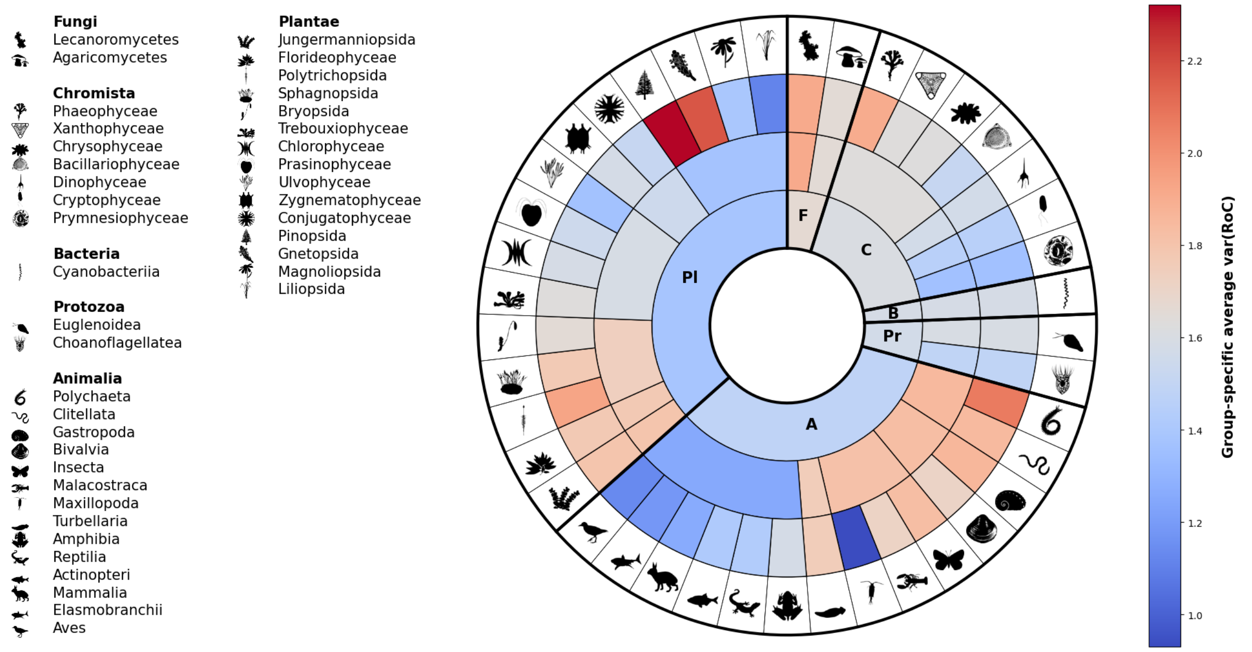


**Figure S9. Association between taxonomy and var(RoC): population-level time series**. For the population-level time series, the nested pie chart shows the average var(RoC) values per taxon (ranks kingdom, phylum, class). Only classes with at least ten populations are shown (41 out of 64 classes: Appendix S1: Figure S3). Each ring stepping towards the center shows a higher taxonomic level. All taxa are ordered clockwise in descending order (classes and phyla within higher taxon). Kingdom abbreviations: F[ungi], C[hromista], B[acteria], Pr[otozoa], A[nimalia], Pl[antae]. Icons: public domain (www.phylopic.org).

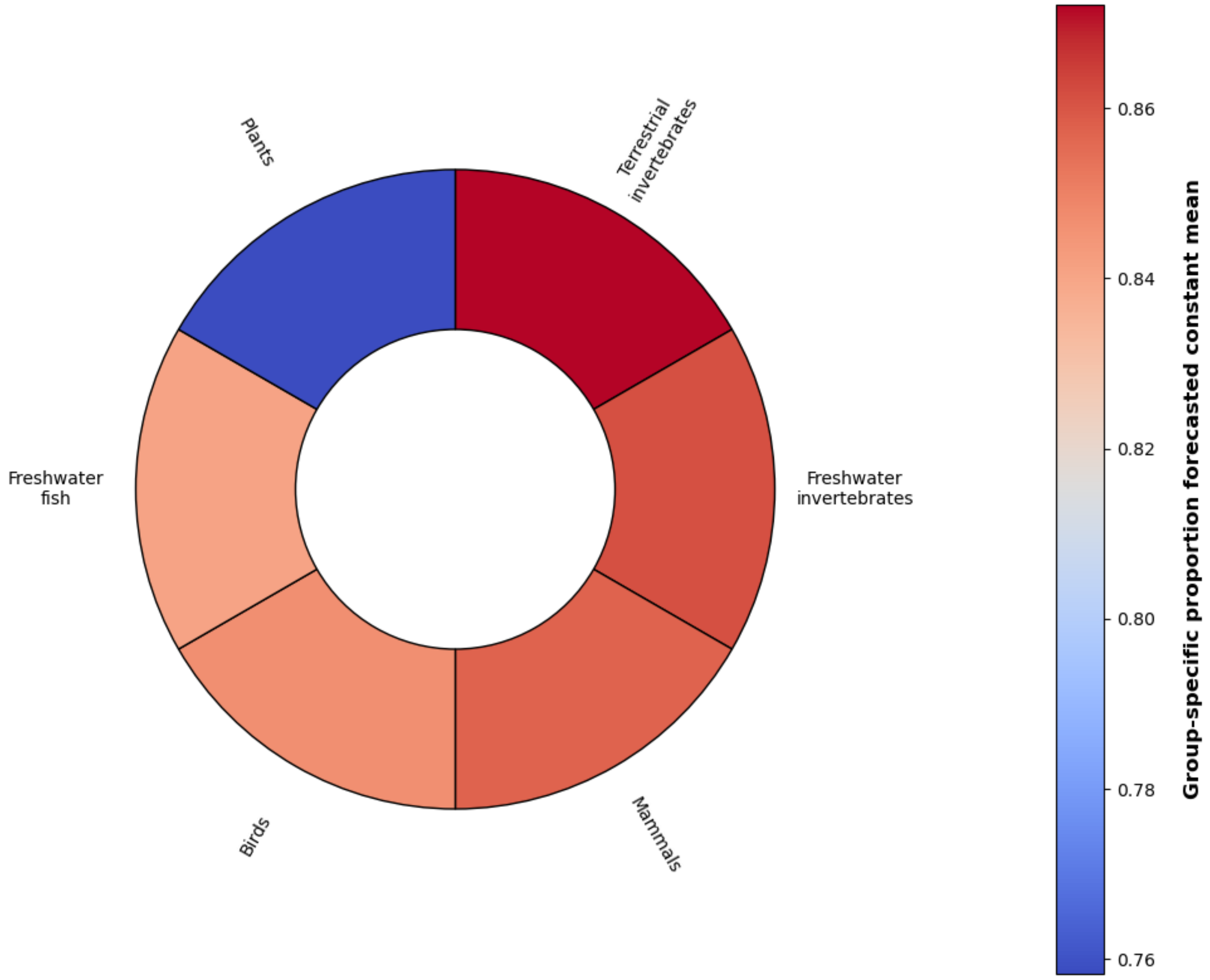


**Figure S10. Association between taxonomy and type of forecasted mean: community-level time series**. Visualization of the taxonomy dependence of the group-level average proportion of communities with forecasted constant mean (colorbar). Groups are shown in descending order.

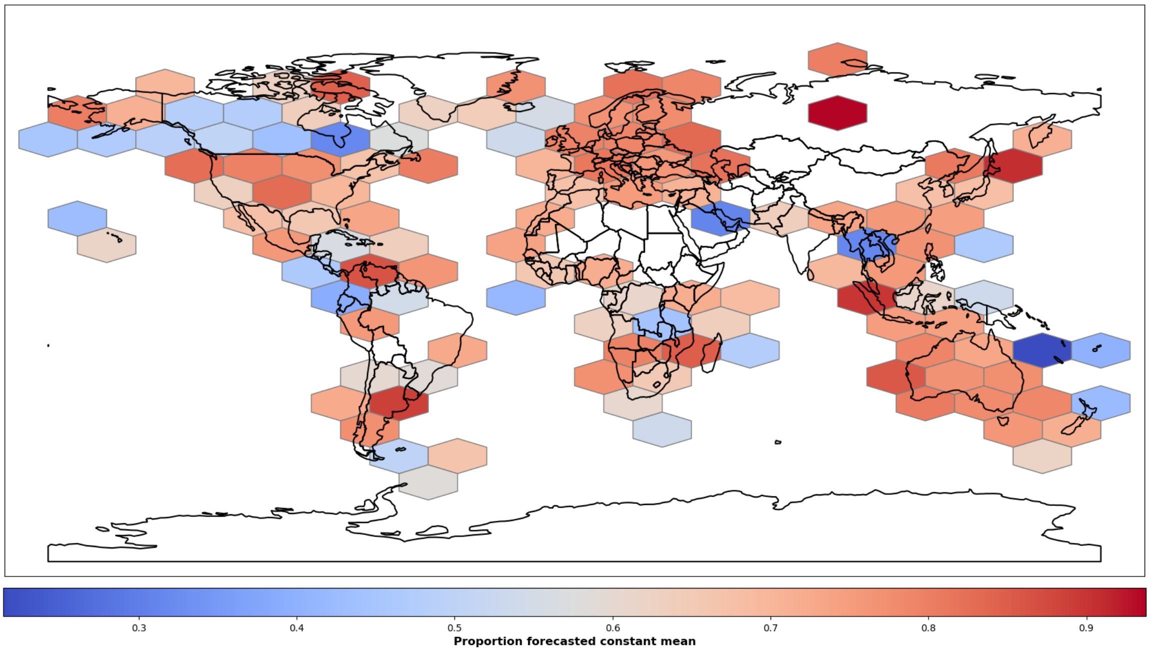


**Figure S11. Association between locality and forecasted constant mean: population-level time series**. The worldmap shows a visualization of the locality dependence of forecasted constant means (colorbar) for the population-level analysis. Only hexagonal bins with at least ten populations are shown (*cf*. Figure S1).

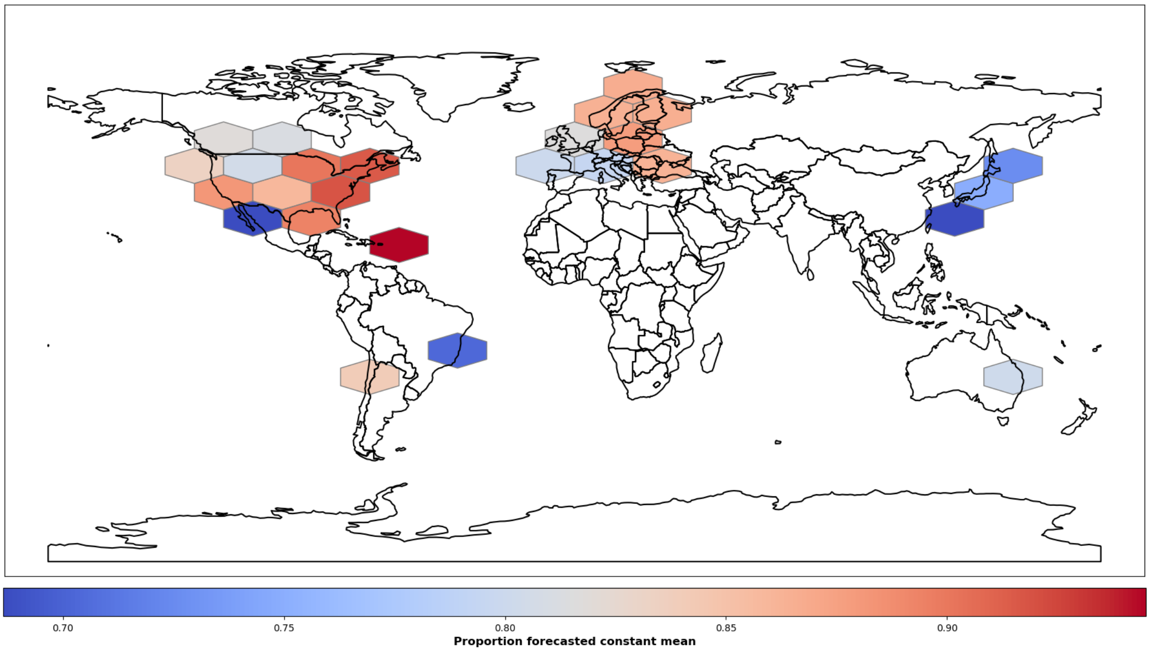


**Figure S12. Association between locality and type of forecasted mean: community-level time series**. The worldmap shows a visualization of the locality dependence of forecasted constant means (colorbar) for the community-level analysis. Only hexagonal bins with at least ten communities are shown (*cf*. Figure S2).

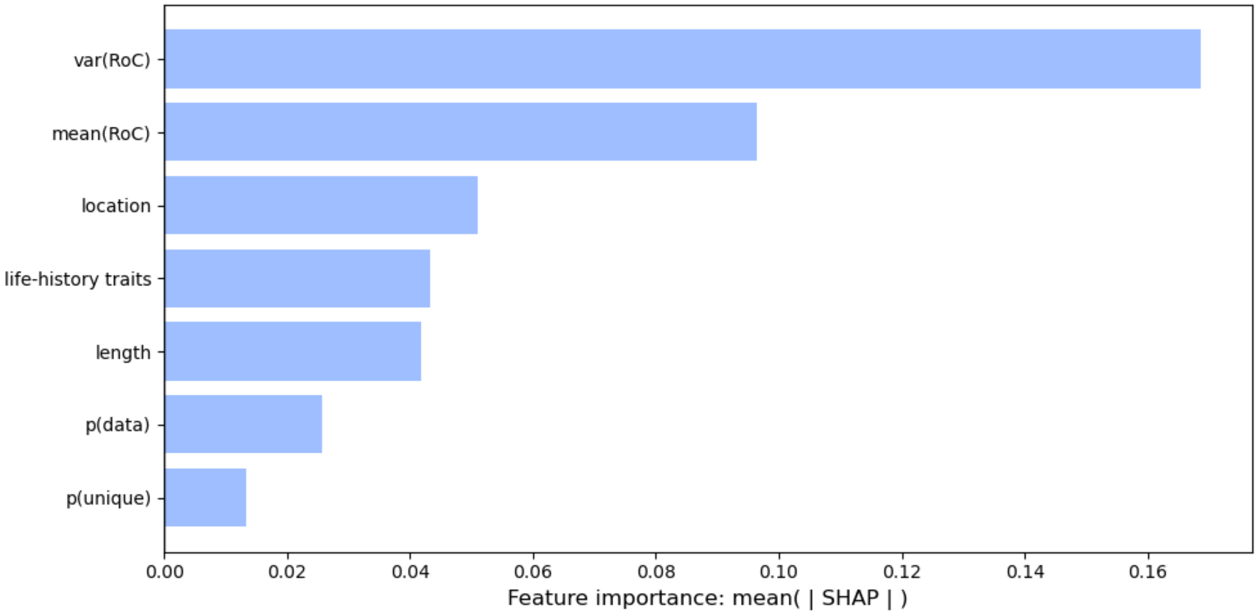


**Figure S13. Feature importance values from the random forest classifiers: amniote populations.** For each feature in the model, the mean absolute SHAP value is shown. Note that 'location' and 'life-history traits' are aggregated (i.e. summed) features: 'location' represents latitude and longitude, and 'life-history traits' comprises the four life-history traits.

#### Supplementary Methods

All references can be found in the reference list of the main text.

**Section S1: Data preprocessing for UCMs**

*Illustrative example: Eastern monarch butterfly population*

We have use the data freely available on https://monarchwatch.org/blog/2025/03/06/monarch-population-status-54/. The time series consists of the yearly total area occupied by monarch colonies at overwintering sites in Mexico. Before fitting the four UCMs, we transformed the raw time-series data as explained in the main text (Materials and Methods: Time-series data).

*Analysis of data collections*

In Table S1 we give a general overview of the time series we used, including online access information. We only analyzed yearly time series; if multiple within-year measurements were available, we used the maximum within-year value. As mentioned in the main text (Materials and Methods: Time-series data), we trimmed every time series so that it started and ended with a non-missing, non-zero value; other inclusion criteria are mention in the main text. Below, we mention specific handlings for some collections.

- BioTIME, macrofungal population-level time series: Because the time series were very short (5 years), we excluded time series where the last value exceeded a count of 1,000. We did not question the reliability of these counts, but compared to the previous years they clearly were 'outliers': such values would have biased trend estimation, given only five years of data, even with UCMs accounting for measurement error.
- GBIF, macrofungal time series: We treated all species-level data from the sampled region as stemming from species-level single populations.
- LTER, macrofungal time series: Because these are presence-absence data, we built yearly counts of presences per species, over the whole sampled region.
- Swiss macrofungal time series: Following the advice of Simon Egli (data owner), we treated species-level counts in the five sampled plots as separate populations.
- Swiss phytoplankton time series: We started all time series with the year 1985. We analyzed 8 sampled sites, located in 7 lakes: Baldeggersee, Greifensee, Hallwilersee, Sempachersee, Vierwaldstättersee, Walensee, Zürichsee (2 sites). See Pomati et al. (2020) for further details on sampling and data.

We fitted the four UCMs (see main text) using the class `UnobservedComponents` from the Python package `statsmodels`.

*Effect of sampling effort in community time series*

Changes in total community abundance over time may reflect changes in sampling effort rather than true biodiversity change, potentially confounding trend results. To assess this, we conducted a proof-of-concept sensitivity analysis restricted to BioTIME communities (Table S1). To this end, we used total reported yearly community abundance as a proxy for sampling effort. For each BioTIME community time series, we fitted the four UCMs to the total abundance time series and applied the same AICc model selection procedure. Community time series where total abundance was best described by a model with a forecasted time trend were temporarily excluded ($n$ = 410). We then recomputed the proportion of α-diversity time series with a forecasted constant mean for the remaining communities. The proportion of forecasted constant means was higher after exclusion (85.3% vs. 81.4%), suggesting our main results are conservative with respect to potential sampling effort confounds.

**Section S2: Feature selection and data preprocessing for random forest classifiers**

*Data preprocessing*

We prepared the results from the UCM fits and the features for the random forest classifiers as follows (separate analysis for population-level and community-level data):

- We only analyzed time series spanning at most 80 years, thus exluding 10 exceptionally long time series. These time series would have unnecessarily stretched the feature's value range (i.e. length, see below), with a very sparse space towards the maximum.
- To ensure all features had a comparable range (to facilitate random forest and SHAP analysis), we applied the following transformations:
  - p(Data), p(Unique): no transformation.
  - length, mean(RoC), var(Roc): transformed to the range 0–1.
  - coordinates: We first transformed these to a unit of kilometer, using the approximation[1] $Latitude' = 111.32 \cdot Latitude,\ Longitude' = 40075 \cdot Longitude \cdot \cos(Latitude \cdot \pi / 180) / 360$. Next, we transformed Longitude' to the range 0–1. For Latitude', we used the range of Longitude' to transform the values to the range 0–1, in order to retain the same overall 'distance' scale.
  - taxonomy: For populations, we first one-hot encoded the categorical taxonomic data. Subsequently, we ran a PCA and selected the first four components, which captured ~78% of the variation in the data. For communities, we only used one-hot encoded taxonomic group information (Table S1).

*Random forest classifiers*

To perform out-of-sample predictions with the random forest classifiers, we used 80% of the respective dataset for training and 20% for testing (predictions). To account for the imbalance between time series with a forecasted constant mean and those with a forecasted time trend (Table 1), we used the `BalancedRandomForestClassifier` from the Python package `Imbalanced learn`, which relies on `scikit-learn`, our main package in this study. The balanced classifier differs from a classical random forest in that it draws a bootstrap sample from the minority class (forecasted time trend) and samples with replacement the same number of samples from the majority class (forecasted constant mean). Finally, for each dataset we performed a 5-fold cross-validation to assess the robustness of the results (Table S2).

[1] As proposed on stackoverflow: https://stackoverflow.com/questions/639695/how-to-convert-latitude-or-longitude-to-meters (accessed March 2025).

*SHAP analyses*

To perform the SHAP analyses, we used the Python package `shap` (Lundgren et al. 2020); specifically, we used the class `TreeExplainer`, suitable for ensemble tree models like random forest classifiers. We used this class to compute SHAP values (Figure S6-S8), and based on these values the overall importance of features (Figure 2) by computing the mean of the absolute SHAP values.

#### Section S3: Investigating drivers of var(RoC)

*Effects of life-history characteristics and location on trend classification (amniotes)*

Based on the findings of Albaladejo-Robles et al. (2023), we investigated the effect of life-history characteristics and locality on trend classification. We restricted the analysis to amniote populations ($n$ = 6916), to match life-history data from Myhrvold et al. (2015). All amniote population data come from the LPI database (Table S1). For taxonomic matching, we only used the species level, thus treating subspecies as species. For location, we used coordinates already used for the main analysis (Section S2). For life-history data, we used the following four attributes: age at maturity, maximum lifespan, body mass, and yearly reproductive output. We computed yearly reproductive output by multiplying the average litter/clutch size and the average number of reproductive events per year. For missing life-history traits in mammals, we supplemented Myhrvold et al. (2015) with data from Pacifici et al. (2013), using age at first reproduction, maximum longevity, and body mass where available. We applied a Box-Cox transformation followed by min-max normalization to all four life-history traits, and a Box-Cox transformation to var(RoC). As for other preprocessing steps and fitting of the model, we followed the approach described in Section S2.

*Relating var(RoC) to process variance (intrinsic ecological dynamics) and measurement variance*

To assess how var(RoC) relates to UCM variance components (eq. 1a-c), we fitted an OLS regression in log-log space. The dependent variable was log(var(RoC) + $\varepsilon$), where $\varepsilon = 10^{-8}$ is a small constant added to avoid log(0). Predictors were (i) log($\sigma_\varepsilon^2$) (measurement variance); (ii) log(signal), where signal means level variance ($\sigma_\xi^2$) or slope variance ($\sigma_\zeta^2$); (iii) an indicator variable for series where process variance is structurally zero (deterministic constant and deterministic trend models). We used HC3-robust standard errors to account for heteroskedasticity. Results are given in Table S4.

We note that var(RoC) and the UCM variance components are derived from the same time series, so the regression does not establish independent attribution of var(RoC) to either source. The elasticity ratio – $\beta$(log(signal)) / $\beta$(log($\sigma_\varepsilon^2$)) – should be interpreted as a descriptive comparison of relative scaling rather than a causal decomposition.
